# Hodgkin and Huxley Opsin Model for Computationally Efficient Optogenetic Neurostimulation in Cells and Networks

**DOI:** 10.1101/2020.11.10.376939

**Authors:** Ruben Schoeters, Thomas Tarnaud, Luc Martens, Wout Joseph, Robrecht Raedt, Emmeric Tanghe

## Abstract

Optogenetics has a lot of potential to become an effective neuromodulative therapy for clinical application. Selecting the correct opsin is crucial to have an optimal optogenetic tool. With computational modeling, the neuronal response to the current dynamics of an opsin can be extensively and systematically tested. Unlike electrical stimulation where the effect is directly defined by the applied field, the stimulation in optogenetics is indirect, depending on the selected opsin’s non-linear kinetics. With the continuous expansion of opsin possibilities, computational studies are difficult due to the need for an accurate model of the selected opsin first. To this end, we propose a Hodgkin-and-Huxley based model (22HH) as alternative to the conventional three and four state Markov models used for opsin modeling. Furthermore, we provide a fitting procedure, which allows for nearly automatic model fitting starting from a vast parameter space. With this procedure, we successfully fitted two distinctive opsins ChR2(H134R) and MerMAID. Both models are able to represent the experimental data with great accuracy and were obtained within an acceptable time frame. This is due to the absence of differential equations in the fitting procedure, with an enormous reduction in computational cost as result. The performance of the proposed model with a fit to ChR2(H134R) was tested, by comparing the neural response in a regular spiking neuron to the response obtained with the non-instantaneous, four state Markov model (4SB), derived by Williams et al. (2013) [1]. Finally, a computational speed gain was observed with the 22HH model in a regular spiking and sparse Pyramidal-Interneuron-Network-Gamma (sPING) network simulation with respect to the 4SB-model, due to the former having two differential equations less. Consequently, the proposed model allows for computationally efficient optogenetic neurostimulation and with the proposed fitting procedure will be valuable for further research in the field of optogenetics.

## I. Introduction

WITH optogenetics, neuronal firing can be controlled with light. This is achieved by genetically expressing opsins, light sensitive ion channels or pumps, in cells or cell subtypes. The merger of this genetic expression and optical stimulation results in superior spatiotemporal resolution with respect to the conventional neuromodulation techniques. Consequently, it is an ideal investigative tool for behavioral studies and a promising biomedical treatment for medical disorders such as epilepsy, Parkinson’s disease and beyond the brain conditions [1]–[7]

The first light sensitive ion channels were discovered in the green alga *Chlamydomonas reinhardtii* by Nagel et al. in 2002 [8]. Of its seven opsin-related genes, two encode light-gated ion channels, i.e., channelrhodopsin-1 (ChR1) and channelrhodopsin-2 (ChR2). These microbial rhodopsins are important for phototaxis and photophobic responses of the alga. Although there exist a 65% homology between them [8], [9], ChR2 is preferred in experiments and applications due to its higher conductance and permeability for cations. Genetic engineering of these has led to a variety of opsins, such as red-shifted, step-function and ultrafast opsins, and mutants with altered ion selectivity [2], [10], [11]. An example of the latter is ChR2(H134R), which is addressed in this paper. This mutation at the inner gate and sodium bindings site, results in an increased *Na*^+^ permeability [12], [13]. Furthermore, other natural versions are being discovered as well, extending the current possibilities even more. An example are the MerMAIDs, which is a family of metagenomically discovered marine anion-conducting and intensely desensitizing channelrhodopsins [14].

In its initial dark-adapted (IDA) state and under voltage clamp conditions, ChR2’s photocurrent exhibits a peak (I_peak_) and a steady-state current (I_ss_) [18]. The peak is reached within 1-2 ms and followed by fast decay onto a steady-state plateau. This is due to light adaptation (Fig. 1 (a, left)). Post-illumination, there is a bi-exponential decay back to baseline, rendering the channel in an apparent dark-adapted state (DA_app_). This is observed by applying a second stimulation after a short period of time (< 10 s), which results in a reduced transient response with a maintained steady-state current (Fig. 1 (a, right)) [18], [19].

**Fig. 1.**
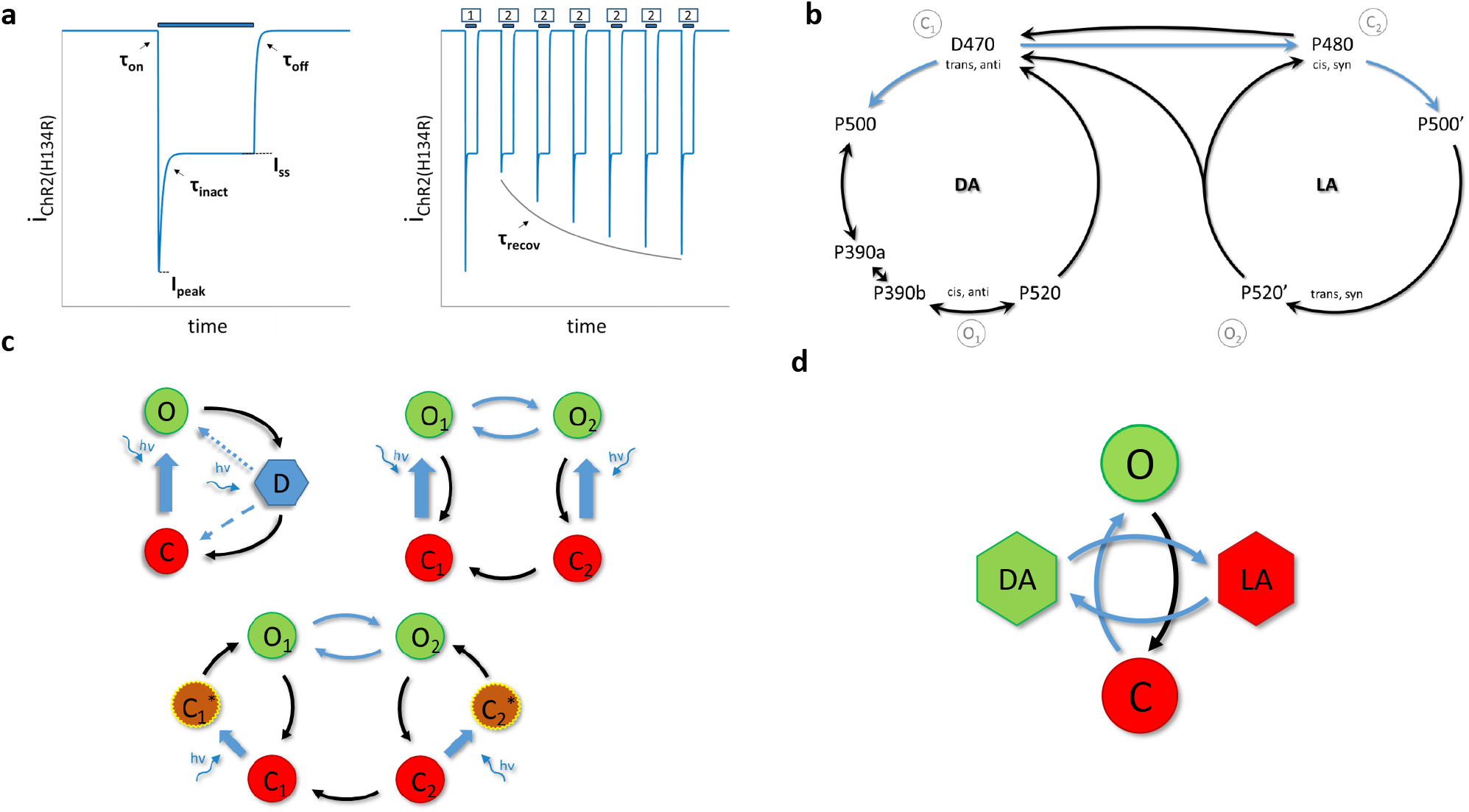
The Channelrhodopsin-2 photocurrent and photocycle. **a**, The photocurrent for a single light pulse on the left. Right, response to a S1-S2 pulse protocol with variable inter-pulse intervals. Light pulses are indicated with blue bars and target features with black arrows. **b**, The unified photocycle based on single-turnover electrophysiology, time-resolved step-scan FTIR, and Raman spectroscopy experiments, as proposed by Kuhne et al. (2019) [15]. **c**, Previously proposed models. (**c**, top left) a three state cycle model with second light dependent step (dotted or dashed step) [16]. (**c**, top right) a four state branching model [1]. (**c**, bottom) a six state model with two extra activation intermediates [17]. **d**, The proposed 2 state-pair Hodgkin-and-Huxley model (22HH). DA and LA indicate dark and light adapted, respectively. O means open, C is closed and D is desensitized.

ChR2 comprises seven transmembrane helices. These are covalently bound with a retinal chromophore forming a protonated retinal Schiff base (RSBH^+^). In its IDA (D470), retinal is in an all-trans configuration [18]. Upon illumination, retinal absorbs photons, rendering it in an excited state. Relaxation triggers a 13 trans-cis isomerization, which initiates a cascade of conformational changes with opening of the pore as result. These spectrally distinct changes are identified with ultraviolet and visible light absorption, and difference infrared spectroscopy measurements (Fig. 1 (b)). The first intermediate (P500) is reached in a picosecond timescale. Next, the RSBH^+^ is deprotonated giving rise to the blue shifted P390 state. This state is in equilibrium with the P520 state, exhibiting a reprotonated RSB. This state is the conducting state. Before returning back to the dark adapted state, the channel converts to a non-conducting state P470. This happens on a millisecond timescale, while complete recovery takes seconds [9], [20], [21].

There is strong evidence for a second photocycle, with similar intermediates. Bamann et al. (2008) [22] identified four kinetic intermediates (P1, P2, P3 and P4) with different lifetimes. Furthermore, light adaptation causes changes in the ion selectivity, i.e., a higher proton and Ca^+^ selectivity for steady-state currents [18], [20], [23]. Also, retinal extraction and Raman measurements indicate a mixture of retinal isoforms occurring in parallel. Two of these, all-trans,15-anti and 13-cis,15-syn retinal, are observed in the DA_app_. As aforementioned, opening of the pore is associated with a retinal trans-cis isomerization. Consequently this gives rise to conducting states 13-cis,15-anti and all-trans,15-syn, respectively. Moreover, electrophysiological measurements identified a bi-exponential, post-illumination current decay and a strong peak-to-steady-state ratio (*I*_ratio_ = *I*_ss_*/I*_peak_) dependence on pH and membrane potential. Neither of this can be explained with a single photocycle model [18], [20], [24]. The transition between the two photocycles is debatable. Either this occurs at the late intermediates (P480 and P480’) or via a photochemical transition at the parent states (D480 and D470). [18], [24]. Recently, Kuhne et al. (2019) [15] proposed an unifying ChR2 photocycle model consisting of two parallel photocycles. They identified three reaction pathways. The classical reaction path is the one described above, with an extra early conducting P390b state. The second path is an early branching to the second photocycle with transition from the D470 directly to P480 due to an all-trans,15-anti to 13-cis,15-syn isomerization. In the third path, the lower conducting P520’ state is reached via a 13-cis,15syn to all-trans,15-syn isomerization [15].

*In silico*, the photocurrent is currently modeled with either a three- or a four-state Markov model (Fig. 1 (c)). This is in accordance with the single and double photocycle hypothesis, respectively. Instead of going through different states before opening as in the UV/Vis-photocycle, the opening is reduced to a single state transition. This is because the D480 → P500 and P500 → P390 transitions occur on a much faster timescale. However, in order to represent fast closure, slow recovery and a steady-state current, a second photon absorption step is proposed for the three state model [13], [20]. The photo-chemical transition either increases the recovery rate or acts as equilibrium modulator between the open and desensitized state. The six state model, as depicted in figure 1 (c, bottom), is an extended version of the four state model. The additional two intermediates are to correctly account for the activation time after retinal isomerizations and to avoid explicit time dependent rates [17]. The four state model is in agreement with the second photocycle hypothesis with modeling of two open and closed states. The transition as depicted in figure 1 (c, top right) is according to the older transition hypothesis, not to the latest unifying photocycle model of Kuhne et al. (2019) [15].

To date there is absolutely no doubt that, with its high cell specificity and temporal resolution, optogenetics wields high potential for neuromodulation tools. Nevertheless, there remain many uncertainties concerning among others, its interference with the intrinsic network dynamics, effects on action potential waveforms and energetic efficiency. *In silico* studies, which allow for extensive and systematical investigation of the effects of the current kinetics are thus invaluable. These studies require an accurate model of the to be investigated opsin. This is then implemented in a biophysically accurate neuron model, which can contain multiple compartments. Furthermore, in case of network studies, multiple of these neuron models are implemented. To date, an accurate model consists of four differential equations [1]. Using such a model therefore increases the computational burden enormously, especially in case of multi-compartment or network studies. Moreover, due to the expanding possibilities, selection of the correct opsin is crucial to have an optimal optogenetic tool. These four state Markov models are not easily fit as they require preliminary knowledge of the parameter space and its complex interactions. Furthermore, finding the optimal parameters is time-consuming as the set of differential equations has to be evaluated at each step in the selected optimization algorithm.

In this study we propose for the first time (to the authors’ knowledge) an alternative for the modeling of the ChR2 photocurrents (Fig. 1 (d)). The proposed model is based on the Hodgkin-and-Huxley model [25], where opening and inactivation are separated. However, instead of inactivation, the second state pair represents the light-dark adaptation, resulting in the accompanied conductance regulation. With this model, the complexity is reduced with fifty percent. Furthermore, by using two light dependent rates in the light-dark adaptation cycle, we hypothesized and realized no loss of ChR2 current features. To this end, a fit is created of the ChR2(H134R) mutant and compared to the 4SB model of Williams et al. (2013) [1]. The performance of both models are tested in a regular spiking neuron [26]. The difference in computation speed is assessed, as well, this in the aforementioned regular spiking neuron for different stimulation patterns and in the sparse Pyramidal-Interneuron-Network-Gamma (sPING) network model [27], [28] with increasing number of transfected neurons. Finally, the versatility of the proposed model is evaluated with a fit to a MerMAID opsin.

## II. Methods and Materials

In this study, a Hodgkin-and-Huxley model with two state pairs (22HH) was tested as alternative for opsin modeling. Below, we first describe the model in full and indicate the link between parameters and certain features. Next, the fitting procedure is elaborated. Finally, we describe the models and metrics used in the analysis of the model performance and computational speed.

### A. The model

The proposed model is based on the original sodium model of Hodgkin and Huxley [25]. It consists of two separate state pairs as depicted in figure 1 (d). In contrast to the sodium model, where the second state pair represents the inactivation gate, the second state pair represents here the light-dark adaptation. This light-dark adaptation of ChR2 affects the conductance of the channel. Therefore, the second state pair acts in this case as a conductance regulator.

After a long enough dark period, the molecules are assumed to be all in closed, dark adapted state. Upon stimulation, the channel opens with a transition C → O. On a slightly slower time scale the equilibrium between dark and light adapted (DA ↔ LA) molecules is reached. The established equilibria of both state pairs depend on the level of optical excitation. After photostimulation, the channels close (O → C). Moreover, they all return to the dark adapted state after a long enough recovery period, which is on a much slower time scale than the other temporal kinetics. Because of this slower time scale, the transition LA → DA has to be light dependent as well. Otherwise the equilibrium would be completely on the side of LA for every optical excitation level. The ChR2 photocurrent can thus be determined as follows:

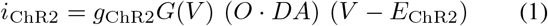

with

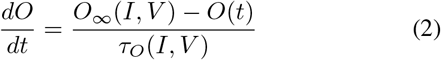

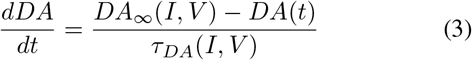

where *g*_ChR2_ is the maximal specific conductivity of the channel, *G*(*V*) is a rectification function, *V* the membrane potential, *I* the light intensity, *E*_ChR2_ the equilibrium potential and *O* the fraction of molecules in the open state, with *O*_∞_ and *τ*_O_ its corresponding equilibrium and time constant. Moreover, *DA* is one when fully dark adapted and (*g*_LA_*/g*_ChR2_) when fully light adapted, with *g*_LA_ the conductivity of a light adapted channel. *DA*_∞_ and *τ*_DA_ are the respective equilibrium and time constants.

Under voltage clamp conditions and a rectangular optical pulse with constant light intensity, the photocurrent can be expressed in a closed form analytical expression:

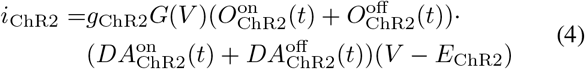

with

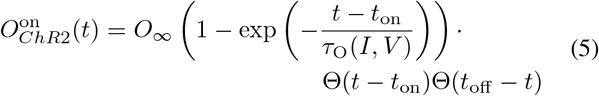

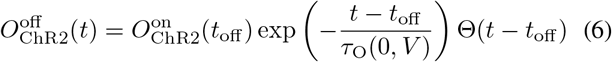

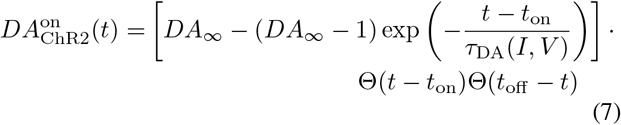

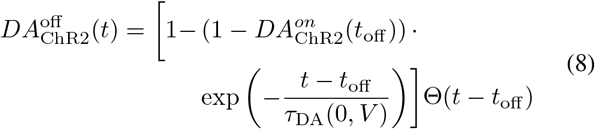

with Θ the Heaviside function and, *t*_on_ and *t*_off_, respectively, the onset and offset of the optical pulse.

Strong correlations between the model time constants and experimentally determined features (Fig.1 (a)) are observed. These can be exploited to obtain a first approximation of the model’s parameters (see section II-B). When *τ*_O_ ≪ *τ*_DA_, the transition rate time constant (*τ*_O_) can be easily obtained from the activation (*τ*_on_) and deactivation (*τ*_off_) time constants. Under the same conditions, *τ*_DA_ strongly correlates with the inactivation time constant (*τ*_inact_) when *I* ≠ 0. The recovery time constant needs to be scaled as shown in eq. 10 to get a good approximation of the dark-light adaptation time constant under dark (*I* = 0) conditions. This relationship is obtained by evaluating the recovery time definition with the given model equations, i.e., *τ*_recov_ = *t*_on,2_ − *t*_off,1_ → *I*_p,2_*/I*_p,1_ = 1 − exp(−1). Here, *t*_on,2_ is the onset time of the second pulse, *t*_off,1_ the offset of the first pulse, and *I*_p,2_ and *I*_p,1_ the current peak value of second and first pulse, respectively.

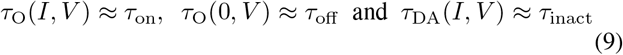

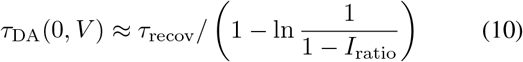

Furthermore, following conditions need to be met for the relationship to hold true:

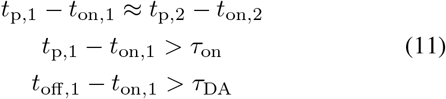

The first, *t*_p,i_ − *t*_on,i_ is the time required to reach the peak value since onset of pulse i. This needs to be approximately the same in both first and second pulse, while these need to be significantly larger than the activation time constant. The last one requires that the steady-state value is reached at the end of the first pulse.

Unless specified, the time constants and time in this study are in seconds, the membrane potential in mV and the intensity in W/m^2^. The units of the conductance depend on the experimental data of each opsin, i.e., mS/cm^2^ and *μ*S in case of the ChR2(H134R) and MerMAID fit, respectively.

### B. The fitting procedure

Due to the dependency on both the potential and light intensity, more than twenty parameters need to be inferred. This vast parameter space impedes finding the optimal solution which is at a high computational cost. To alleviate this, the fitting procedure can be divided into four steps.

The first step is the extraction of the features, which is described by Williams et al. (2013) [1]. The peak current (*I*_peak_) is the maximal deflection from baseline. The steady-state current (*I*_ss_) is the plateau value. The current ratio (*I*_ratio_) is then *I*_ss_*/I*_peak_. The time constants are extracted using mono-exponential curve fits. To this end, a nonlinear least-squares curve fit is performed, with a trust-region-reflective algorithm. Furthermore, a multi-start algorithm with ten starting points was used to ensure finding of the global solution. The variable and function tolerance were set to 1e-12. The recovery time constant, i.e., the time necessary between two pulses to have a second peak current which is 63 % of the first peak (see definition previous subsection), was determined from a set of two-pulse experiments.

Next, *τ*_O_ and *τ*_DA_ are fit to the obtained target data. Both are fit to the corresponding time constants (see eq. 9 and 10) using the aforementioned nonlinear least-squares method. Again, a multi-start algorithm is used but with 2000 starting points. For the intensity dependence, sigmoidal functions on the log-scale are used while for the voltage dependence a logistics model was selected. The two dependencies are combined by either a multiplication or a reciprocal addition. The relationships and combination schemes are given by equations 12–16, with p_i_, i = 1 → 6 indicating the unknown parameters of each relationship individually.

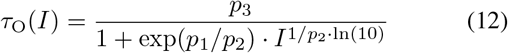

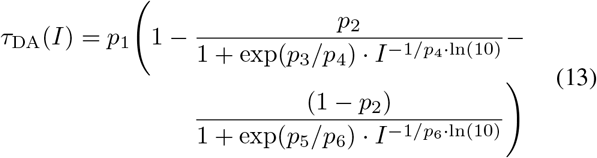

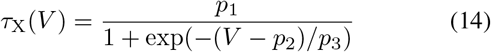

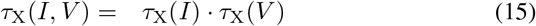

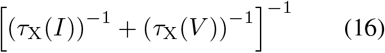

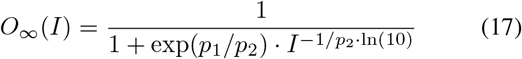

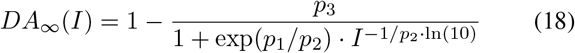

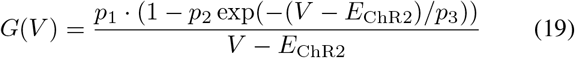

In a third step, the parameters of the rectification function *G*(*V*) and the equilibrium constants *O*_∞_ and *DA*_∞_ are fit. The used relationships are given in equations 19, 17 and 18, respectively. The potential dependence of *O*_∞_ and *DA*_∞_ are omitted because this is mostly covered by the rectification function. The parameter values are determined by minimizing the cost function described below:

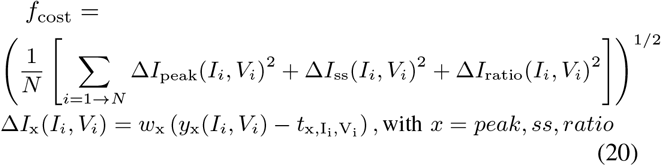

Here, *y*_x_ and 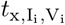 are respectively the model output and target value at stimulation values (*I*,*V*), with 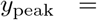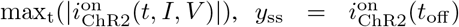 and 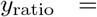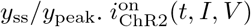 is the current during the photostimulation pulse (*t ϵ E* [*t*_on_, *t*_off_]) for a certain intensity *I* and voltage *V*. The current is calculated by evaluating the equations 4 - 8 with the determined dependencies in the previous step. *N* is the total number of stimulation sets (*I*,*V*). The minimization of *f*_cost_ is performed with the MATLAB *fmincon*-function and multi-start algorithm with 3000 starting points to increase chance of finding the global optimum. The upper and lower boundaries as well as the initial conditions are summarized in table I. Extra nonlinear constraints are applied to assure that *O*_∞_ approaches one for high intensities (see section III-A and III-D) and *G*(*V*) ≥ 0. A final constraint ensures a current decay back to baseline after the optical stimulation, i.e., *i*_on_(*t*_off_) > *i*_off_ (*t*) or *O*^on^(*t*_off_) · *DA*^on^(*t*_off_) > *O*^off^ · (*t*) *DA*^off^ (*t*), resulting in:

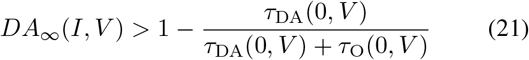

**TABLE I.** Summary of all parameters. The boundary conditions, lower bounds (LB) and upper bounds (UB), of first 2 steps of the fitting procedure and initial values (X0). the intermediate (-intm) and final (-final) model parameters of the selected mutant and model, with RSRS the H134R mutant with a double reciprocal addition combination of time constant dependencies (*τ*_*X*_ (*I*) and *τ*_*X*_ (*V*), EQ. 16), PP the H134R mutant with a double product combination of time constants dependencies (EQ. 15) and the MerMAID fit (MM). Parameters that varied between ChR2(H134R) and MM fit are separated with vertical line. Between brackets is another parameter solution, resulting in the same model.

| | $\tau_O(I)$ | | | $\tau_O(V)$ | | | $\tau_{DA}(I)$ | | | | | | $\tau_{DA}(V)$ | | |
| --- | --- | --- | --- | --- | --- | --- | --- | --- | --- | --- | --- | --- | --- | --- | --- |
|  | P1 | P2 | P3 | P1 | P2 | P3 | P1 | P2 | P3 | P4 | P5 | P6 | P1 | P2 | P3 |
| LB | -10 | 0 | 0 | 0 | -100 | -1000 | 0 | 0 | -10 | 0 | -10 | 0 | 0 | -100 | -1000 |
| UB | 10 | 20 | 1 | 100 | 100 | 1000 | 10 | 1 | 10 | 20 | 10 | 20 | 100 | 100 | 1000 |
| X0 | 1 | 1 | 0.5 | 1 | -50 | 10 | 1 | 0.5 | 0 | 0.125 | 3 | 0.5 | 1 | -50 | 10 |
| RSRS-intm. | 1.93 | 0.68 | 0.022 | 23.26 | 0.14 | 12.40 | 10 | 0.56 | -1.59 | 0.88 | 1.96 | 0.11 | 100 | -38.94 | 14.70 |
| RSRS-final | 1.81 | 1.17 | 0.021 | 23.14 | -0.39 | 13.19 | 10 | 0.56 | -1.58 | 0.87 | 1.96 | 0.11 | 99.74 | -38.69 | 12.02 |
| PP-intm. | 1.99 | 0.67 | 0.034 | 0.64 | -89.16 | 14.31 | 6.74 | 0.50 | 2.00 | 0.11 | -1.3 | 0.88 | 1.50 | -70.01 | 19.13 |
| PP-final | 1.93 | 0.88 | 0.030 | 0.63 | -88.67 | 8.37 | 6.73 | 0.50 | 1.98 | 0.11 | -1.28 | 0.88 | 1.66 | -64.54 | 28.55 |
| MM-intm. | 1.70 | 1.49 | 0.035 | 0.29 | 48.56 | 738.24 | 0.17 | 0.0081 | -2.80 | 14.69 | 1.05 | 0.42 | 24.77 | 80.92 | 164.75 |
| MM-final | 3.70 | 3.35 | 0.037 | 0.20 | 49.99 | 718.60 | 0.18 | 0.0082 | -3.00 | 15.57 | 0.998 | 0.429 | 24.42 | 80.87 | 172.82 |

| | $O_\infty(I)$ | | $DA_\infty(I)$ | | | $G(V)$ | | | $g_{Chr2}$ | $E_{Chr2}$ |
| --- | --- | --- | --- | --- | --- | --- | --- | --- | --- | --- |
|  | P1 | P2 | P1 | P2 | P3 | P1 | P2 | P3 |  |  |
| LB | -10 | 0 | -10 | 0 | 0 \| 0.8 | 0 \| - | 1.1 \| - | 0 \| - | - \| 0 | - \| -100 |
| UB | 10 | 20 | 10 | 20 | 1 | 100 \| - | 100 \| - | 500 \| - | - \| 100 | - \| 100 |
| X0 | 1 | 1 | 1 | 1 | 0.1 \| 0.9 | 1 \| - | 10 \| - | 50 \| - | - \| 30 | - \| 0 |
| RSRS-intm. | 3.45 | 0.71 | 2.03 | 0.13 | 0.71 | 9.91 (1) | 1.24 | 46.17 | 1 (9.91) | 0 |
| RSRS-final | 3.38 | 0.62 | 1.96 | 0.12 | 0.77 | 10.77 (1) | 1.25 | 44.52 | 1 (10.77) | 0 |
| PP-intm. | 3.45 | 0.71 | 1.99 | 0.15 | 0.73 | 8.93 (1) | 1.27 | 42.37 | 1 (8.93) | 0 |
| PP-final | 3.44 | 0.68 | 2.25 | 0.065 | 0.75 | 9.10 (1) | 1.27 | 41.47 | 1 (9.10) | 0 |
| MM-intm | 3.76 | 0.40 | 0.74 | 0.52 | 1.00 | - | - | - | 62.00 | -3.64 |
| MM-final | 3.67 | 0.39 | 0.40 | 0.54 | 0.9987 | - | - | - | 62.22 | -3.62 |

Finally, a global optimization is performed with the parameters of all rate functions included. First, a new parameter space is defined, which is 10% of the original parameter space but centered around the values obtained in previous steps and limited by the former. With the gathered dependencies, the ChR2 current is calculated according to equation 4. All model features are now extracted in the same manner as performed on the experimental data. These are used to determine a cost function which is the weighted root mean square error (see equation 20), with additional terms: Δ*τ*_on_(*I, V*)^2^, Δ*τ*_off_ (*I, V*)^2^, Δ*τ*_inact_(*I, V*)^2^ and Δ*τ*_rec_(*I, V*)^2^. Subsequently, the problem is optimized with a bounded particle swarm optimization [29]–[31], containing 1000 particles and with a time limit of 24 hours. The same solver settings and constraints are imposed as described in previous steps. The single-pulse experiments are evaluated with a time step of 1.5e^−4^ s, while for the two-pulse experiments a step of 1 ms is used.

### C. Performance tests

In this study, two opsin fits were performed. First, a fit is made to the data reported by Williams et al. (2013) of the ChR2(H134R) [1]. The model accuracy is compared to the four state Markov model created by the same group. Four metrics are used to analyze the goodness-of-fit, i.e. Root mean square error (RMSE), Root mean square normalized error (RMSNE), Root mean square weighted error (RMSWE) and root mean square Z-score error (RMSZE):

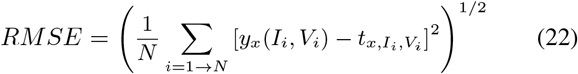

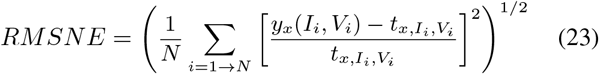

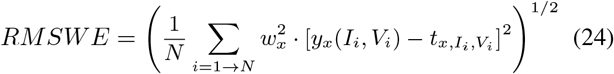

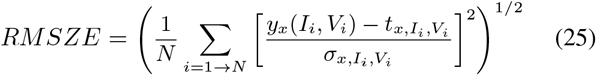

with *y*_x_(*I_i_, V_i_*), 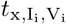 and 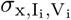 respectively the model output, target feature and standard deviation of target feature x under intensity *I* and voltage *V* of set i, and *w*_x_ the weights used in *f*_cost_. The metrics are also determined in the overall, time constant features (*τ*_on_ + *τ*_off_ + *τ*_inact_ + *τ*_rec_) only and current features (*I*_p_ + *I*_ss_ + *I*_ratio_) only case. Here the squared errors of all features are summed first before taking the root and mean. The RMSWE is equivalent to the training error. However, it could not be used to compare the model fits as the used weights were not equal across fitting procedures (different weights were used in the 4SB fit, see Williams et al. (2013) [1]). Therefore, the other metrics were defined as well. Where the RMSE is biased by high values, the RMSNE is biased by values close to zero and RMSZSE could not be determined for the recovery time constant.

Both models are then implemented in a regular spiking neuron, described in Pospischil et al. (2008) [26]. The strength duration curves (SDC) are determined. When the irradiance is selected as strength for the SDC, a poor fit is obtained. This is due to the assumption of an RC equivalent circuit and a rectangular stimulation pulse in the Hill-Lapicque relationship (eq. 26) [32], [33]. Therefore, the SDC fit is performed on the average inward stimulation current or temporal averaged current (*i*_ChR2,avg_, TAC), as described by Williams and Entcheva (2015) [32].

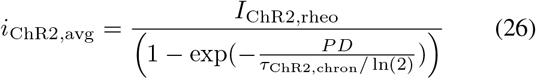

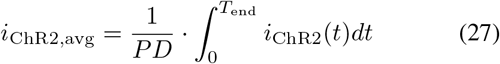

with PD the pulse duration and *T*_end_ one second after the end of the pulse. The relationship between the irradiance and *i*_ChR2,avg_ is obtained through a power series fit, which allows calculation of the irradiance rheobase (*I*_rheo_) and chronaxie (*τ*_chron_) as follows:

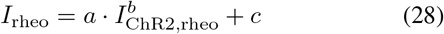

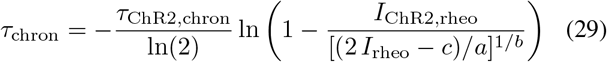

where a, b and c are parameters obtained in an empirically power series fit of the irradiance curve versus the inward stimulation current (*I* = *a* · (*i*_ChR2,avg_)^*b*^ + *c*) [32].

Moreover, the simulation speed is determined for different stimulation paradigms, i.e., simulation time (*T*_end_)/runtime in a regular spiking neuron [26]. Therefore, we varied the pulse repetition frequency, stimulation time and duty cycle. The intensity was fixed for each model and set to a value that elicited a firing rate of 100 Hz in the regular spiking neuron in case of a two pulse stimulation of two seconds with duty cycle 0.5 and pulse repetition frequency of 1 Hz. The models were solved by the MATLAB Variable Step Variable Order solver (VSVO) ode113-solver (order 1-13, Adams-Bashort-Moulton predictor-corrector pairs) [34], [35], with a maximum time step of 100 *μ*s and default tolerances, i.e., relative and absolute tolerance equal to 1*e*^−3^ and 1*e*^−6^, respectively.

Finally, computational gain with the proposed model compared to the 4 state Markov model was tested in a network model with an increasing number of transfected neurons. Therefore, we used the sparse Pyramidal-Interneuron-Network-Gamma (sPING) [27], which was implemented via the DynaSim toolbox [28]. The ChR2(H134R) models were added to the pyramidal neurons. The number of inhibitory neurons was varied between 3 and 100 while the 4/1, pyramidal/interneuron-ratio was maintained. The network was fully connected and the GABAa and AMPA conductivities were scaled such that the total input per neuron stayed the same, i.e., *g*_GABAa_ = 2/(*N*_intern_)[mS/cm^2^] = *g*_AMPA_, with *N*_intern_ the number of interneurons in the sPING-network. In each case a single pulse stimulation of 300 ms was applied with a total simulation time of 500 ms. The irradiance was set such that the firing rates were equal for both ChR2 models. The study was performed with both a fixed step (10 *μ*s) runge-kutta 4 solver and an ode15s-solver (stiff VSVO-solver, order 1-5, based on numerical differentiation formulas) [34], [35] with a maximum time step of 100 *μ*s, and a relative and absolute tolerance of 1e-6.

The results shown in this paper are computed with a 3.4 GHz clock rate, quad core system and 8 GB RAM.

### D. Versatility

The versatility of the proposed model is shown with a fit to the MerMAID1 opsin [14]. Photocurrent traces were obtained in HEK-293 cells during a 500 ms illumination with 500 nm light. Recordings were made between −80 and +40 mV with a step of 20 mV at an intensity of 3734 W/m^2^. A light titration at −60 mV with intensities between 0.46 and 3734 W/m^2^ was performed as well. The two-pulse experiment was only performed at −60 mV and with 3734 W/m^2^ illumination. For more detail we refer to the work of Opperman et al. (2019) [14]. The same metrics as aforementioned are used to assess the fit accuracy.

## III. Results

To test the feasibility of the proposed Hodgkin-and-Huxley model with two state pairs (22HH) in opsin modeling, it was fit to two data sets. First, we fitted the model to the data set of a ChR2(H134R) opsin reported by Williams et al. (2013), which was collected in a ChR2(H134R)-HEK293 stable cell line [1]. By the same group already a four state Markov model was fit. This allowed us to analyze the performance of our model in detail. To this end, a comparison of the response to optical stimuli was made in a regular spiking neuron [26]. Moreover, the computational speed was determined for different stimulation paradigms in the former neuron model as well as in the sPING [27] network model with increasing number of transfected neurons. Finally the versatility of the proposed modeling scheme was assessed with a fit to a MerMAID opsin which is an anion-conducting and intensely desensitizing channelrhodopsin.

### A. The ChR2(H134R) fit

A 22HH fit of the ChR2(H134R) opsin was obtained by applying the fitting procedure, described in the materials and methods section II-B, to the experimental data. As Williams et al. (2013) [1] already reported the target features, the first step could be omitted. The absence of differential equations in our fitting procedure allowed for multiple fits to be made, due to the significant reduction of the computational cost. Multiple weight sets, non-linear constraints and combinations of dependency addition of the time constants (product eq. 15 and reciprocal sum eq. 16) were tested. The parameters of the two best fits are shown in table I, where RSRS and PP is the fit with a double reciprocal sum and product combination, respectively. Both results were obtained with *w*_peak_ = 10, *w*_ss_ = 20, *w*_ratio_ = 50, *w*_on_ = 1000, *w*_inact_ = 1000, *w*_off_ = 1000, *w*_recov_ = 20, and a constraint where *O*_∞_(*I, V*) > 0.6 for *I* ≥ 5500 W/m^2^. The weights are chosen as such to level the differences between features to the same order of magnitude. As a result, all features have the same impact in the cost function with a slight preference for the current features. The time constant features are all expressed in seconds, while their values are in the order of milliseconds (except *τ*_recov_), explaining the high weight values. The extra constraint is justified as the current peak already starts to saturate for the highest intensity values, thus clamping the intensity dependence of the open steady-state value above the bending point in the logistics curve.

The models’ accuracy according to the four goodness-of-fit metrics (eq. 22–25) are shown in figure 2. Overall, a positive effect of the final optimization step can be observed. The largest impact is on the time constants, as expected. In the second step of the fitting procedure, the transition rate time-constants (*τ*_O_ and *τ*_DA_) are approximated with a one on one relationship of the target features (see eq 9 and 10). These approximations are true in case of high differences in order of magnitude. However, when the differences are smaller some cross correlations exist, for instance *τ*_DA_ strongly affects *τ*_on_ as well, resulting in a underestimation of *τ*_O_. We denote that according to all metrics, the estimation accuracy of *τ*_on_ and *τ*_inact_ increases, however, at the cost of *τ*_off_. Also, a significant improvement is observed in case of *τ*_recov_. This deviation is due to the fact that the conditions (see eq. 11) are not fully met. Furthermore, an increased goodness-of-fit of the inactivation time constant can be observed in case of the RSRS vs PP fit. *τ*_DA_ predominately defines both the inactivation and recovery time constant. In case of the PP fit, a separation of variables is applied where independence is assumed. However, as can be seen in figure 3 (f) and (h), a more clear voltage dependency is present in *τ*_recov_ compared to *τ*_inact_. In other words, for low intensities, with high time constants as result, the potential effect is high while the effect is low for high intensities or small time constants. This interdependence is exactly obtained with the reciprocal addition scheme. The same, however less pronounced, can be observed in case of the activation and deactivation time constants (*τ*_on_ and *τ*_off_). Consequently, only the RSRS fit is used in further analysis.

**Fig. 2.**
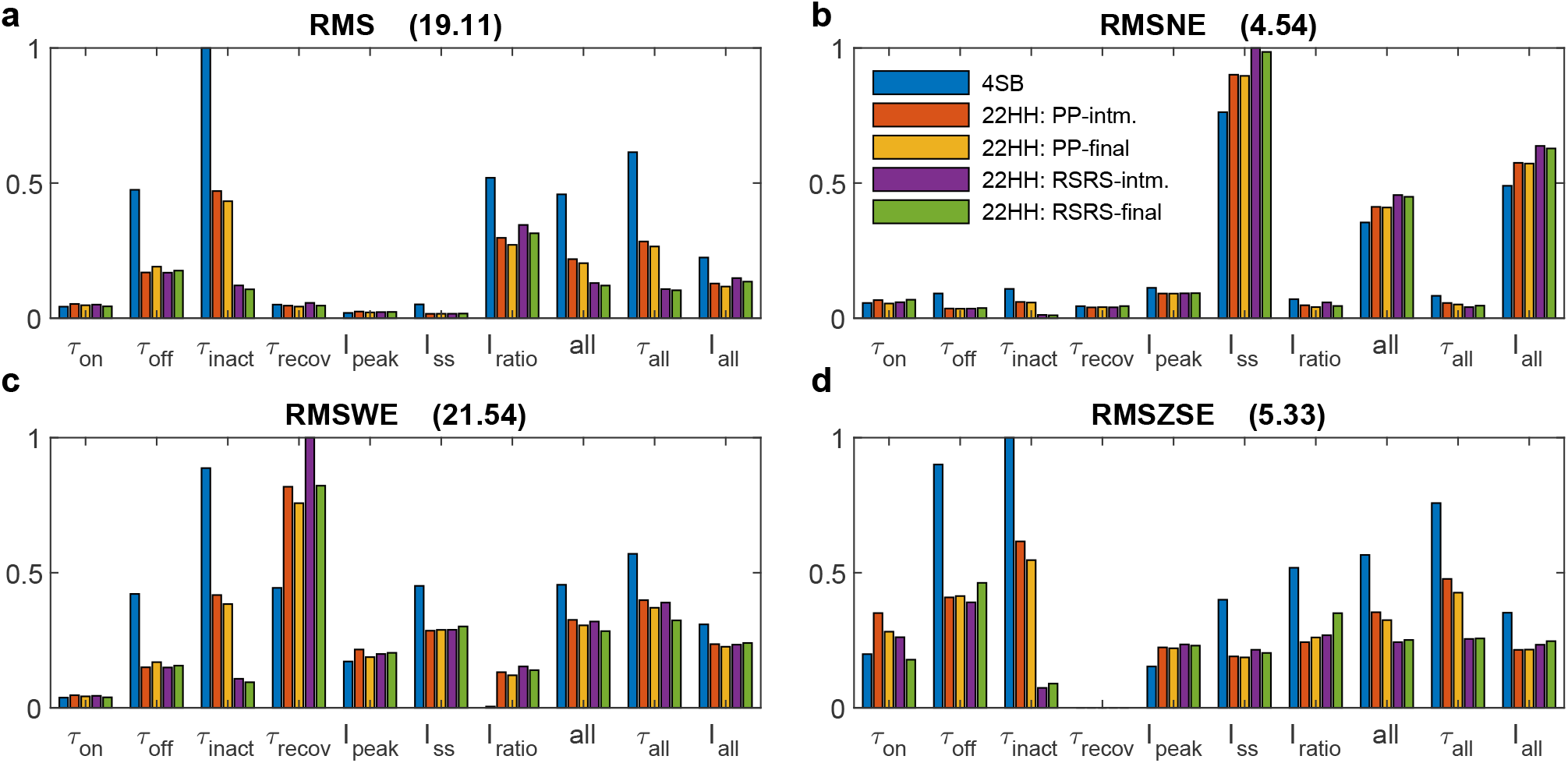
Normalized goodness-of-fit results of the model fits to the ChR2(H134R) data reported by Williams et al. (2013) [1]. Goodness-of-fit according to four metrics: Root-mean-square error (eq. 22, **a**), Root-mean-square normalized error (eq. 23, **b**), Root-mean-square weighted error (eq. 24, **c**) and Root-mean-square of z-score error (eq. 25, **d**). The compared models are the 4SB model reported by Williams et al. (2013) [1], the 22HH model with twice the product combination of time constants (intermediate fit 22HH: PP-intm and final fit 22HH: PP-final), and the 22HH model with twice the reciprocal addition combination of time constants (intermediate fit 22HH: RSRS-intm and final fit 22HH: RSRS-final). The normalization value is given in the title of each sub-figure. *all*, *τ*_all_ and *I*_all_, are errors where the squared errors of all features, all time constants and all current features are, respectively, added first before taking the root and mean. The depicted legend is valid in all sub-figures

**Fig. 3.**
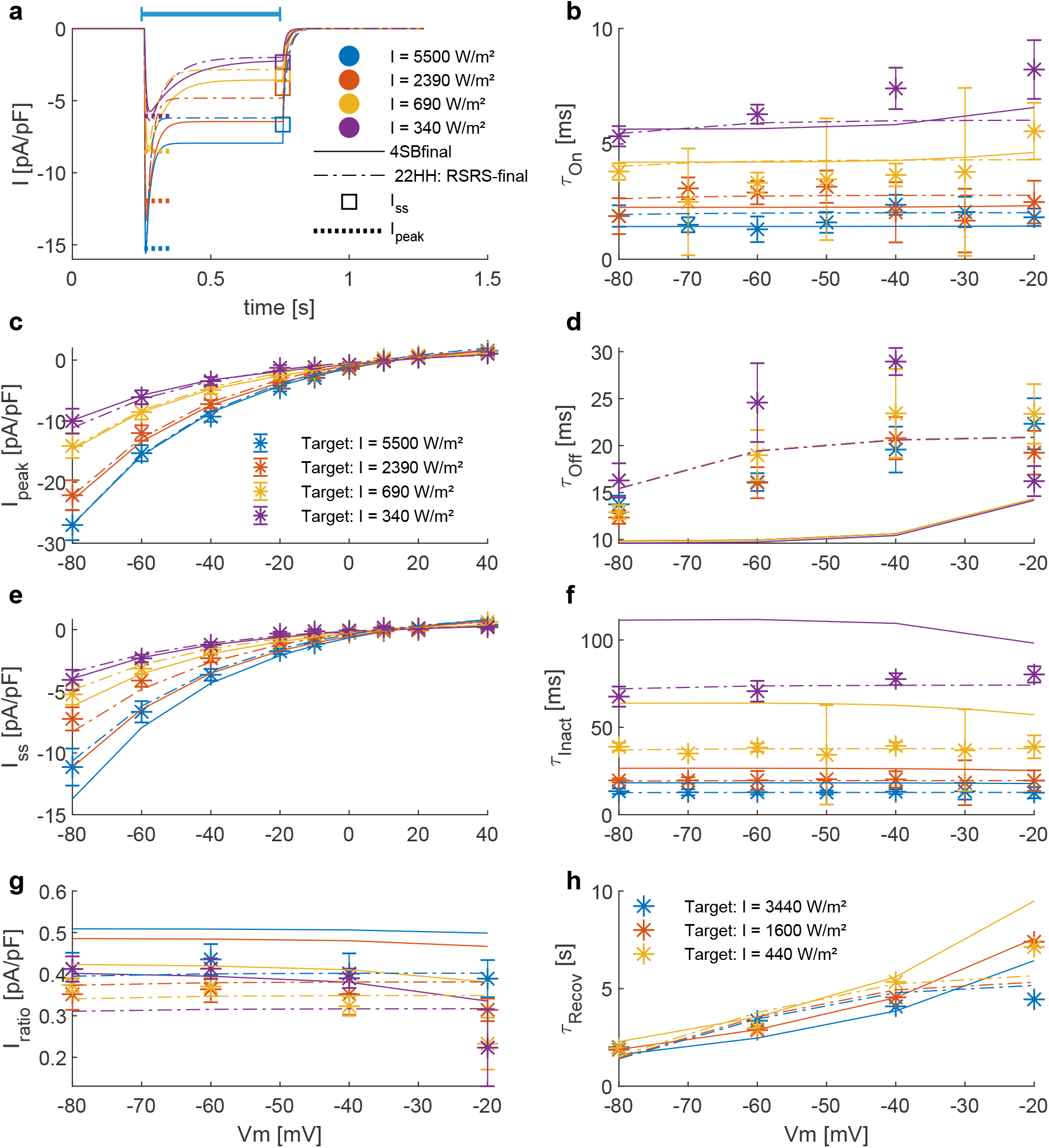
Comparison of model outcomes (4SB and 22HH: RSRS-final) with parameters obtained from experiments. **a**, The ChR2(H134R) current during a pulse of 0.5 s (indicated by blue bar) at a voltage clamp of −60 mV; according to the 4SB model (full lines) and 22HH model (dashed dotted lines). The colors indicate the applied intensity and are valid in **a-g**. The dotted line and square indicate respectively the experimental current peak and steady-state current at corresponding intensity and potential. **b,d and f**, Voltage dependence of respective *τ*_on_, *τ*_off_ and *τ*_inact_ across four irradiance levels. **c,e and g**, The current-voltage curves of the peak, steady-state and current ratio, respectively. The asterisks with errorbars indicate the experimental mean ± standard deviation. **h**, The recovery time constant as function of the membrane potential for three different irradiance levels as depicted in the plot.

Figure 3 shows a detailed comparison of the outcome of our model according to the RSRS fit and the 4SB model, versus the experimentally determined target features. Overall, it can be observed that the proposed model performs at least as well as the 4SB model. Moreover, all features are well approximated. It can be seen that with the 4SB model, the steady-state value is overestimated in case of negative potentials (Fig. 3 (a) and (e)). However, a better representation is obtained for positive potentials, which explains the lower root-mean-squared normalized error (RMSNE, Fig. 2 (b)).

### B. Neural response in regular spiking neuron

To analyze the neural response, the strength duration curves (SDC) are determined of the proposed 22HH model with RSRS fit and the 4SB model in a regular spiking neuron, described in Pospischil et al. (2008) [26]. First, the Hill-Lapicque model fit is performed on temporal average current (TAC), as described in section II-C. Very good fits were obtained for both models. The adjusted r^2^ 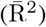 of TAC versus PD are 0.9961 and 0.9953 for the 22HH and 4SB model, respectively. The rheobase of the 22HH model (0.49 *μ*A/cm^2^) is slightly higher than when the 4SB is used (0.47 *μ*A/cm^2^). Also the chronaxie is higher (47.51 ms vs. 39.45 ms). Consequently, according to the 4SB model for any pulse duration, less charge is injected optogenetically to excite a regular spiking neuron via a ChR2(H134R) opsin. The difference between the models can be attributed to the difference in deactivation time constant (*τ*_off_). This is higher in the 22HH model resulting in a slower closing mechanism and thus increased current injection after the AP. A good cell-type-specific empirical mapping of TAC to irradiance was obtained as well (eq. 28), with 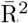 values of 0.9449 (22HH) and 0.9638 (4SB). The parameter values are respectively, a = 8.18, b = 1.26 and c = 1.6798, and a = 22.30, b = 1.51 and c = 12.32 in case of the 22HH and 4SB mapping. The lower 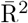 of the 22HH mapping resulted also in a slightly lower value of 0.9298 for the irradiance to PD curve while this is 0.9509 in case of the 4SB fit. Based on the mapping parameters and figure 4, it can be seen that lower intensity level results in higher injected currents when the 22HH model is used. Indeed, extrapolation of the model fit to low intensities results in higher open probabilities than for the 4SB model, hence the difference in irradiance rheobase of 4.90 W/m^2^ versus 19.01 W/m^2^. Based on the higher peak values for high intensities in case of the 4SB model, one could expect convergence of the irradiance SDCs. However, at small pulse durations and due to the slow activation kinetics, the peak value is not reached. Even though the activation time constant is overall higher for the 22HH model (Fig. 3 (b)), the bi-exponential current rise due to the extra state variable (*τ*_ChR2_ · *dp/dt* = *S*0(*I*) − *p*, a time-dependent function reflecting the probabilistic, non-instantaneous response of the ChR2-retinal complex to light [1] in the 4SB model results in a lower current value at the end of the pulse.

**Fig. 4.**
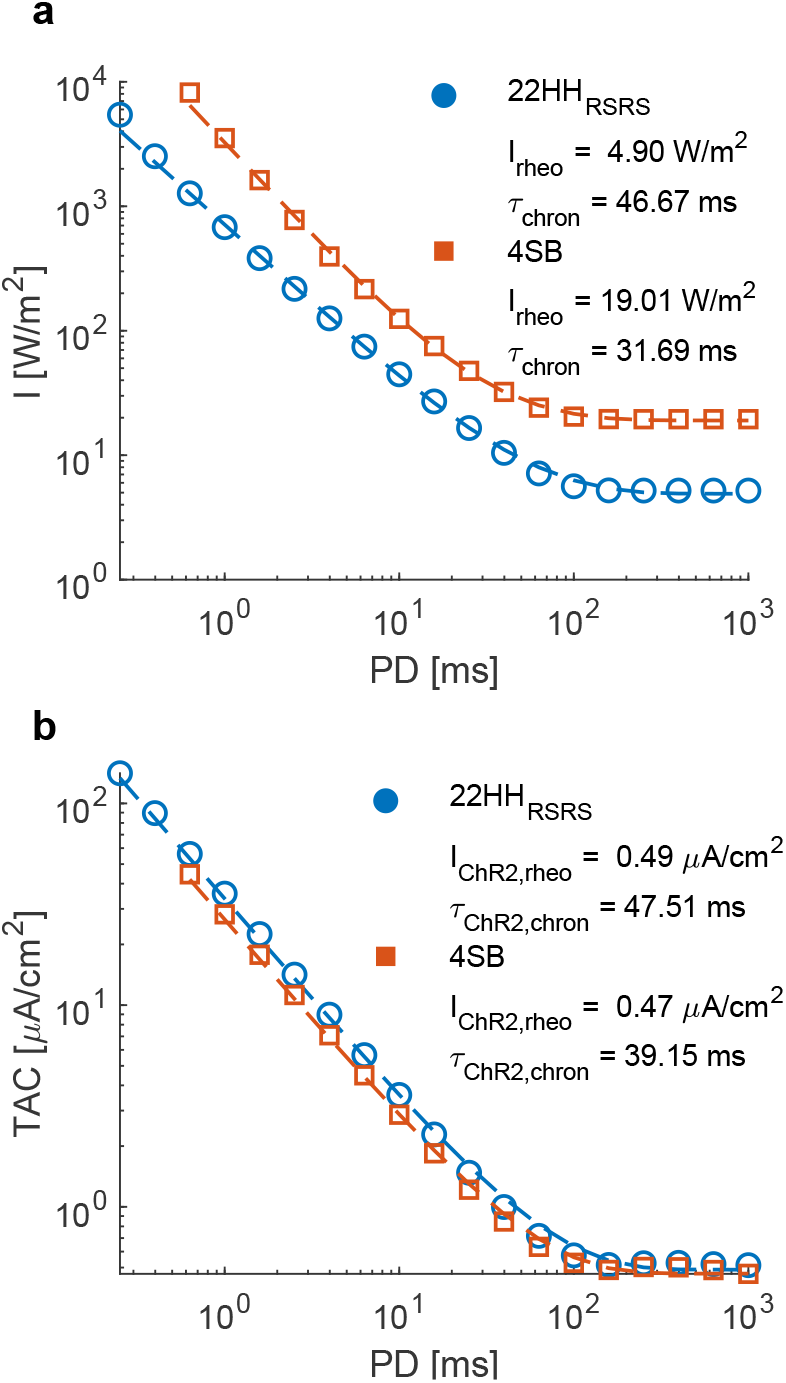
The strength duration curves (SDC) of the 22HH RSRS and 4SB model in a regular spiking neuron. **a**, Irradiance versus pulse duration with a mapping (dashed line) of the SDC in **b** according to a power series. **b**, Temporal average current or average injected current vs pulse duration. Dashed line represents the Hill-Lapicque model fit. The rheobase and chronaxie are depicted in the figures. The results of the 22HH and 4SB model are in blue and red, respectively

### C. Computational speed

The proposed model in this study contains only two differential equations, which is 50% less in comparison with the 4SB model. Consequently, a reduction of the computational time is expected. Figure 5 (a-f) summarizes the computational speed for different stimulation protocols in a regular spiking neuron. This for fixed irradiances (22HH: 3162 W/m^2^ and 4SB: 1259 W/m^2^) set to a value that elicit a firing rate of 100 Hz, as described in section II-C. Subfigures 5 (a-d) show an overall increase of the computational speed in favor of the 22HH model, with a maximum of 25% for high frequency and duty cycle stimulation. On average the relative difference of the simulation speeds, i.e., simulation speed with 22HH minus simulation speed with 4SB with respect to the latter, is about 20%. Because the simulations were solved using a variable step solver, the difference in firing rate could distort the effective simulation speed, as during an action potential a smaller timestep is selected. Therefore, the relative difference of the simulation speed normalized to the firing rate is depicted as well, with an increase of the gain to 60% as result. The runtime versus number of transfected neurons is depicted in figure 5 (g-i). The simulation outcomes were the same with the variable and fixed step solver, validating the solver settings. Moreover, the firing rate was equal for both opsin models, hence no normalization was necessary. A clear reduction can be observed when the 22HH model is selected instead of the 4SB model, both with a fixed and variable step solver. The time gain by using the proposed model is 15% (5%) in case of 12 neurons and goes up to 40% (15%) and rising when 400 transfected neurons are included with a variable (fixed) step solver.

**Fig. 5.**
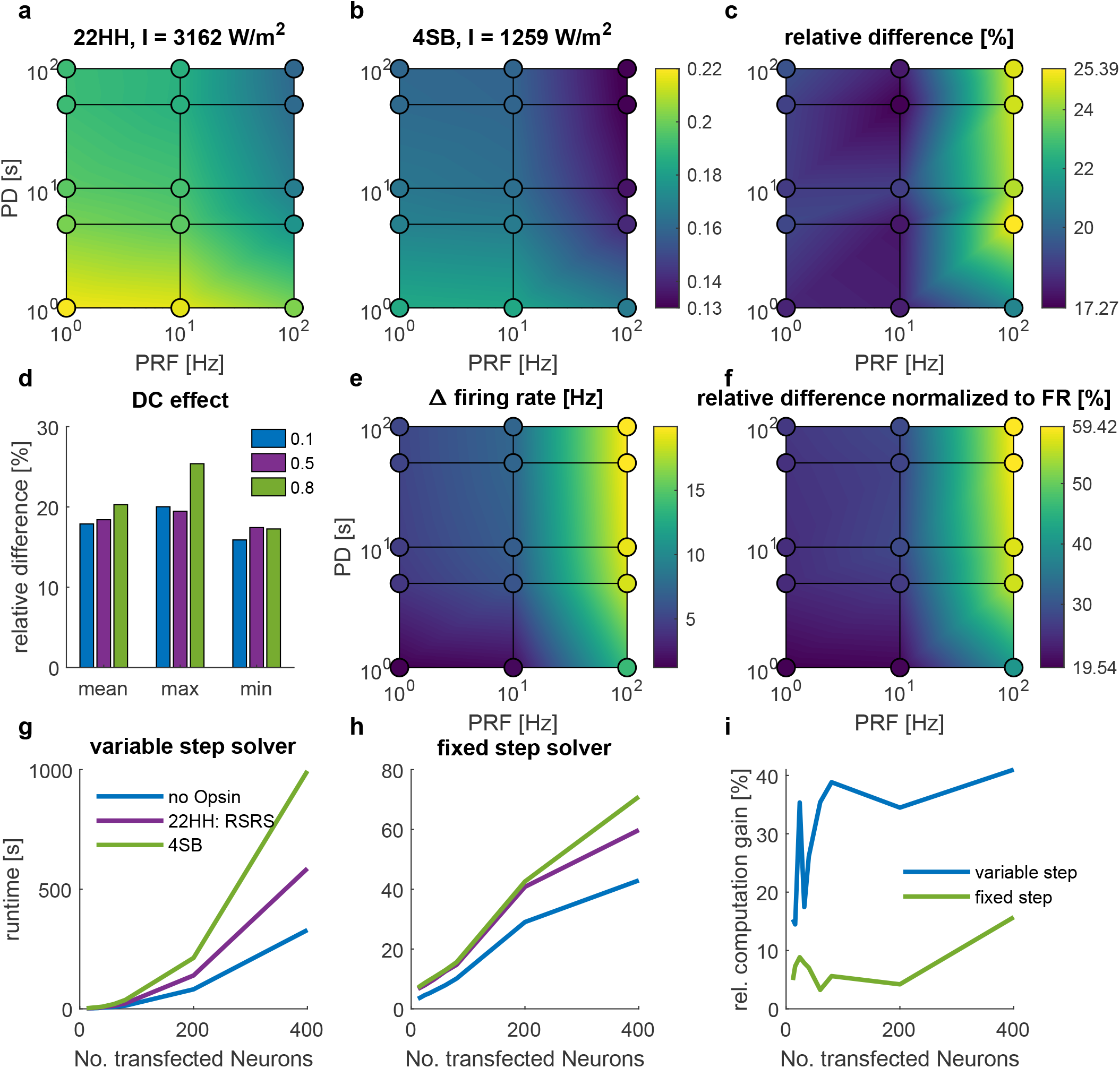
The computational speed of optogenetic neuromodulation in a regular spiking (RS) neuron and sparse Pyramidal-Interneuron-Network-Gamma (sPING). **a-f**, Simulation speed, i.e. simulation time/runtime for different stimulation protocols with varying pulse duration (PD) and pulse repetition frequency (PRF) in a regular spiking neuron, described by Pospischil et al. (2008) [26]. **a**, The absolute simulation speed with the 22HH-RSRS fit. **b**, The simulation speed with the 4SB model. Colorbar is valid for **a** and **b**. **c**, The relative difference, i.e., (22HH-4SB)/4SB. **d**, The effect of the duty cycle on the simulation speed. **e**, The difference in firing rate in case of the 22HH model vs. 4SB. **f**, The relative difference of simulation speed normalized to the firing rate. **g-i**, Runtime of a continuous 300 ms optical pulse in the sparse Pyramidal-Interneuron-Network-Gamma (sPING), with increasing number of transfected neurons. **g**, Runtime with a variable step solver. **h**, Runtime with a fixed step solver. **i**, Relative computation gain, i.e., -(22HH-4SB)/4SB. The used intensities are shown in the titles of **a** and **b**, which give rise to a 100 Hz firing rate (see section II-C)

### D. Versatility of the proposed model

Finally, we address the versatility of the proposed model and the fitting procedure. Due to the increasing number of possible opsins, it is favorable that their kinetics can be correctly modeled and a fit is easily obtained without preliminary knowledge. To this end, we applied our fitting procedure to experimental data of a MerMAID opsin, which has unlike classical ChR2 a very strong desensitization [14]. Starting from the photocurrent traces, the target features had to be extracted first. Next the parameter space was defined. The rectification function was omitted because this was not observed in the experimental data. Aside from this, the lower bound and initial condition of only the third parameter of the dark adaptation cycle was altered (Table I). This straight forward adjustment was made due to the strong desensitization. The weights of the cost function were set to *w*_peak_ = 0.04, *w*_ss_ = 1, *w*_ratio_ = 250, *w*_on_ = 10000, *w*_inact_ = 10000, *w*_off_ = 10000, *w*_recov_ = 10, again to level the errors to the same order of magnitude. Because no saturation of the current was observed at high intensity levels a constraint: *O*_∞_(*I, V*) ≤ 0.50 for *I* ≤ 4000W/m^2^, was added.

The result of the fit is shown in figure 6. The parameters of the final and intermediate fit are summarized in table I. The model here is with a double product combination of the time constant dependencies. Because the recovery time constant was only determined under one condition, there is no evidence on the interdependence of the variables. This is also supported by the small voltage dependence of the (de)activation time constants. Overall, it can be stated that a good fit is obtained as all kinetics are expressed correctly. Only, the deactivation time constant seems to be underestimated. However, this is due to the trade off between this and the steady-state current value to ensure a current decay back to baseline after the optical stimulation, as denoted in Methods and Materials II-B and equation 21.

**Fig. 6.**
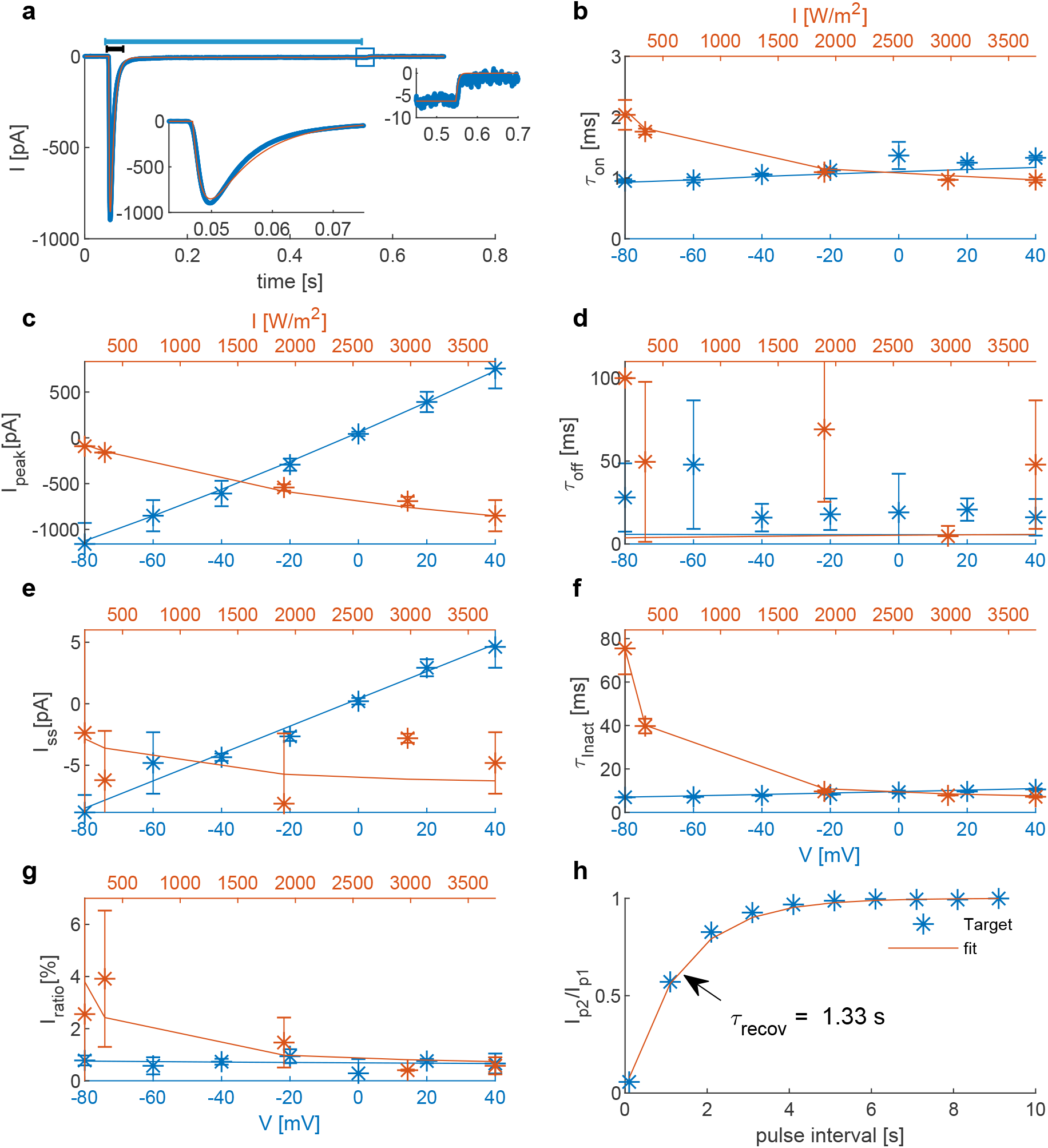
Comparison of the 22HH-Mermaid (final fit) model outcomes and experimental data. **a**, In blue, the photocurrent of a voltage clamp experiment during a 0.5 s continuous illumination (indicated with blue bar on top) of 500 nm light with an intensity of 3734 W/m^2^ [14]; In red, corresponding model outcome. Left inset is zoom of the current peak (0.045-0.075 s, indicated with black bar). Right inset is zoom of current deactivation (0.45-0.7 s, indicated with square). **b-g**, The voltage dependence of the target features (*τ*_on_, *I*_peak_, *τ*_off_, *I*_ss_, *τ*_inact_ and *I*_ratio_) in blue at an irradiance of 3734.4 W/m^2^ and light dependence in red at a holding potential of −60 mV. **h**, Ratio of the peak currents in response to a two-pulse stimulation protocol at −60 mV and 3734 W/m^2^ as function of the inter-pulse interval. The recovery time (the interval time necessary to have a ratio of 63%), is indicated with a black arrow.

## IV. Discussion

The proposed Hodgkin-and-Huxley type model for the modeling of opsins appears to be a good alternative to the computationally more expensive four state Markov, non-instantaneous models. All features are represented, with even some improved fit accuracy in comparison with a four state Markov variant. Furthermore, with the proposed fitting procedure, we were able to fit two opsins, ChR2(H134R) and MerMAID. Although the prominent difference of the mutants kinetics, the fitting procedure allowed us to get these fits with only minor adjustments of the parameter space and constraints. Therefore, creating the possibility for automatic model fitting based on photocurrent traces. Moreover, a good fit is obtained within an acceptable time frame, due to the absence of differential equations in the fitting procedure. The intermediate fit is obtained within three hours, while the final fit always flagged the time limit of 24 hours. Increasing the limit improves the fit accuracy but only small changes were observed. Fine tuning of the optimization settings, such as number of particles or tolerances, could reduce the training error even more. However, this is out of the scope of this study.

The proposed model is an empirical model. The fit is performed on a limited dataset thus extrapolation should be treated with care. This is clear from the neural response results in section III-B. Although both the 4SB and our model were fit to the same experimental data, a clear discrepancy between the fitted rheobase is observed 4.90 W/m^2^ (22HH) versus 19.01 W/m^2^ (4SB). Unlike the chronaxie where the difference can be attributed to the model’s structure, the difference in rheobase is due to the discrepancy between opening rates after extrapolation to low intensities, attributed to the fit and intensity dependence chosen in each model. More experiments are required in order to validate this.

The dependencies chosen here are all, except the rectification, sigmoidal. Therefore, they are all bounded and monotonic. This is in accordance with a channel’s behavior, i.e., increased and faster opening at higher intensities but limited to an open probability of one. We opted for a biphasic logistics function for *τ*_DA_(*I*) modeling. This is in agreement with the hypothesis of the necessity of two light dependent rates (DA → LA) and (LA → DA), see section II-A and the second and third photochemical pathways described by Kuhne et al. (2019) [15] (Fig. 1). Other functions were tested, e.g., weibull or asymmetric logistics with double intensity dependence, however no improvement was observed. Initially, separation of variables was assumed to suffice due to the lack of experimental evidence of complex channel interdependence of both irradiance and potential of each feature separately. However, due to the models structure, *τ*_on_ and *τ*_off_ share the same voltage dependency, as well as *τ*_inact_ an *τ*_recov_. The voltage dependence of *τ*_recov_ and *τ*_off_ was clearly more pronounced in the experimental data of the ChR2(H134R) mutant. Therefore, the reciprocal addition equation 16 was tested as alternative, resulting in an improved fit accuracy. However, this only scales down the voltage dependent effect on *τ*_on_ and *τ*_inact_ while the same relationship is maintained. The necessity of more complex relationships could be investigated in future work as well as the need for voltage dependence of the rate functions steady-state values (*O*_∞_ and *DA*_∞_), which was omitted in this study.

Currently the model incorporates voltage and irradiance dependence. Studies have however shown the importance of pH on the channel kinetics in many opsins. Furthermore, ion concentrations have an impact on the reversal potentials and current rectification [20], [36]. Schneider et al. (2013) [23], postulated a model based on the kinetics of multiple ion species interacting with the channel, with an improved representation of the current rectification [1], [23]. While the photocurrent properties are unaffected by pH-changes, the MerMAID photocurrent is strongly dependent on the Cl^−^ concentrations. The fit performed here was on experimental data recorded with an extracellular Cl^−^ concentration of 150 mM and intracellular Cl^−^ of 120 mM, explaining the depolarizing currents (negative sign in fig. 6) as an anion conducting channel. By changing the extracellular concentration to 10 mM^1^, the channel’s reversal potential is shifted to the reversal potential of Cl^*−*^. Evidence of the Cl^−^ effect on channel kinetics is still absent but further experiments are needed [14]. Consequently, the model fit shown here can be used in computational studies but the reversal potential should be adjusted accordingly.

With the current model structure, the model responds instantaneously to light. With the 4SB model this is circumvented by adding a extra state variable with a time constant of 1.5 ms. It is clear that for long (PD ≫ *τ*_on_) continuous pulses its effect is negligible, as activation is dominated by the activation time constant. However, with short bursts or pulses, this non-instantaneous activation becomes prominent as observed in section III-B. In future work, it could therefore be interesting to incorporate this non-instantaneous response. This could probably be obtained by adding an extra state variable, as performed with the 4SB model, however at the cost of the computational speed. Another possibility is to raise the open state, O(t), to a higher power, smoothing the transition but without irradiance control. Modification of the model’s structure could be circumvented by gradually increasing the intensity, instead of applying a rectangular pulse.

## V. Conclusion

To facilitate computational studies in the field of optogenetics, we proposed a Hodgkin-and-Huxley based model as alternative to the conventional three and four state Markov models. In the proposed model, the second state pair acts as a conductance regulator, modeling the light-dark adaptation cycle. With this model type, a reduction in complexity is obtained resulting in only two differential equations compared to four in case of the preferred, non-instantaneous four state Markov models used for opsin modeling. With the provided fitting procedure, nearly automatic model fits of two distinctive opsins ChR2(H134R) and MerMAID were obtained. Both model fits were performed within an acceptable time frame thanks to the absence of differential equations and parameter space reduction associated with the multi step approach. Moreover, both models are able to represent the experimental data with great accuracy. Due to the model’s structure, there is, however, an instantaneous response to light, overestimating the injected current at very short pulses. Furthermore, pH and ion concentration dependence are not incorporated. In its current state with only two differential equations, the computational speed is increased up to 25% in a regular spiking neuron and up to 40% in a network of 400 transfected neurons.

## ACKNOWLEDGMENT

The authors would like to thank Johannes Oppermann for sharing his experimental data on MerMAIDs. R. Schoeters is a PhD Fellow of the FWO-V (FR) (Research Foundation Flanders, Belgium). T. Tarnaud is a PhD Fellow of the FWO-V (SB) (Research Foundation Flanders, Belgium).

Exploration of the parameter space and dependencies in this work was carried out using the Supercomputer Infrastructure (STEVIN) at Ghent University, funded by Ghent University, the Flemish Supercomputer Center (VSC), the Hercules Foundation and the Flemish Government department EWI.

## Footnotes

1 The concentrations are exchanged with respect to a conventional neuron, where the typical intracellular and extracellular concentrations are 10 mM and 120 mM, respectively. This explains the experimentally measured depolarizing currents (negative sign), while one would expect hyperpolarizing currents (positive sign) from a Cl^−^ conducting channel.

## Notes

### Competing Interest Statement

The authors have declared no competing interest.

